## Supplementary figures and images for "Cryptic Sex in *Leishmania* Depends on *SPO11* Paralogs"

### Figure S1

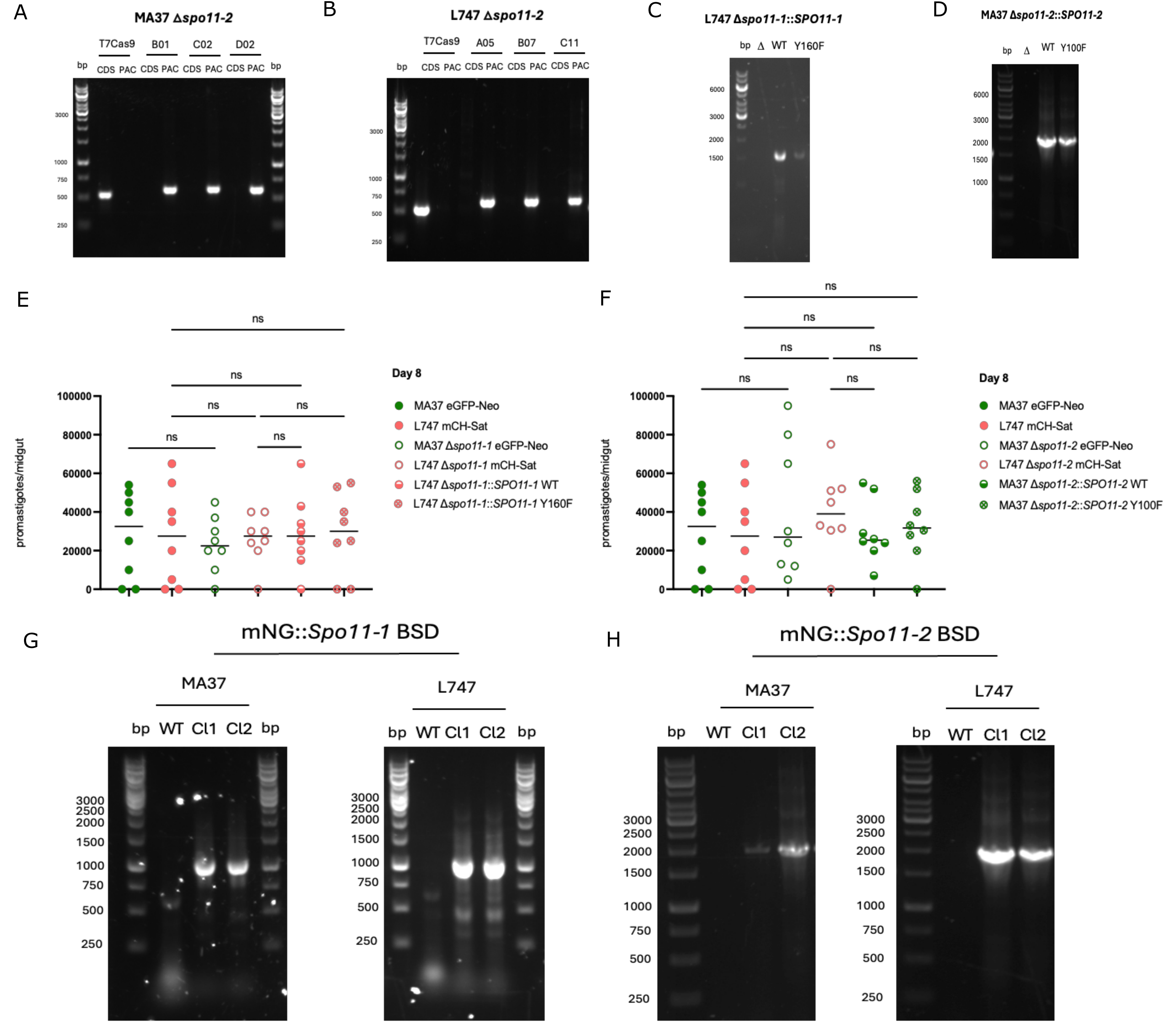

### Figure S2

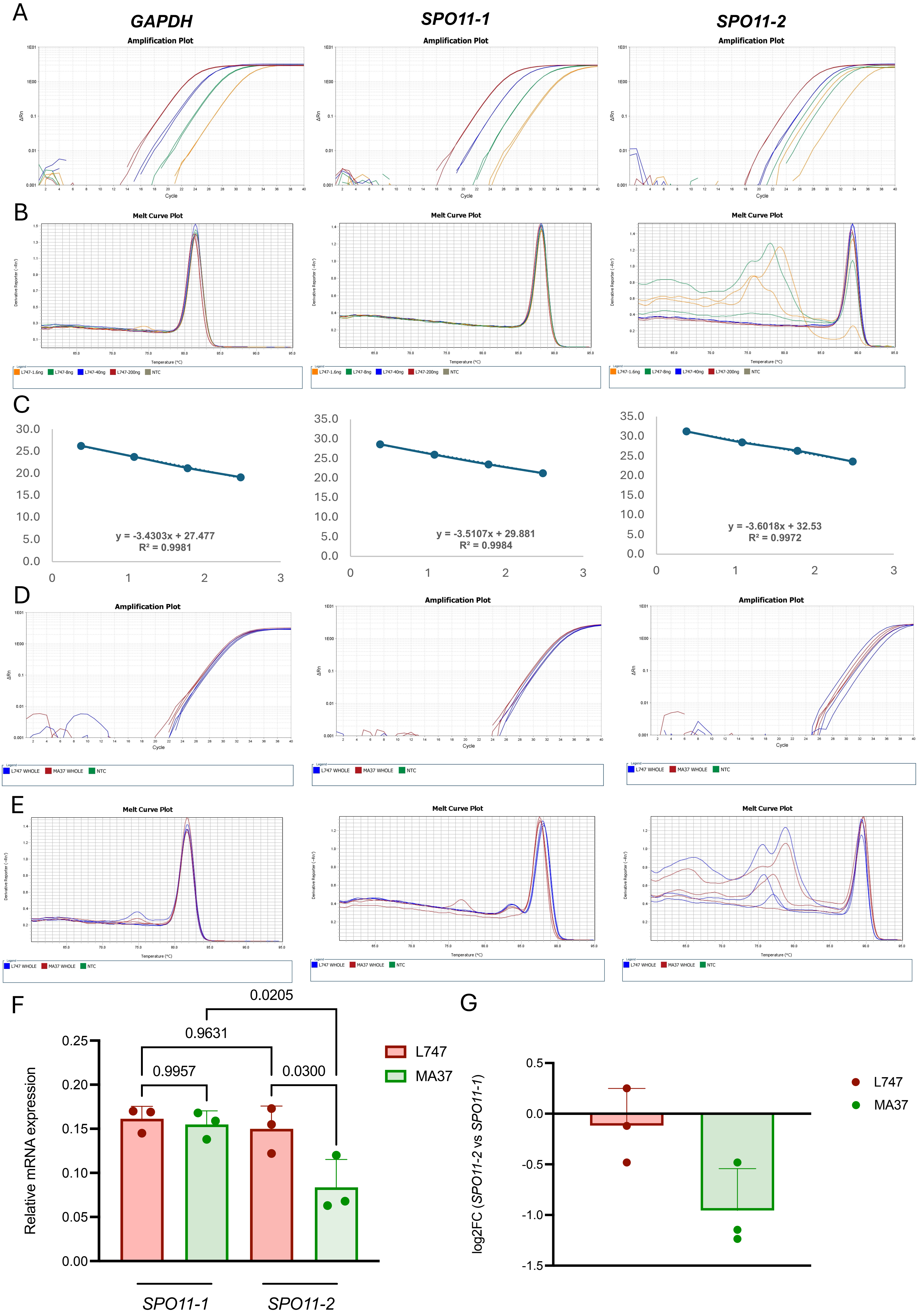

### Figure S3

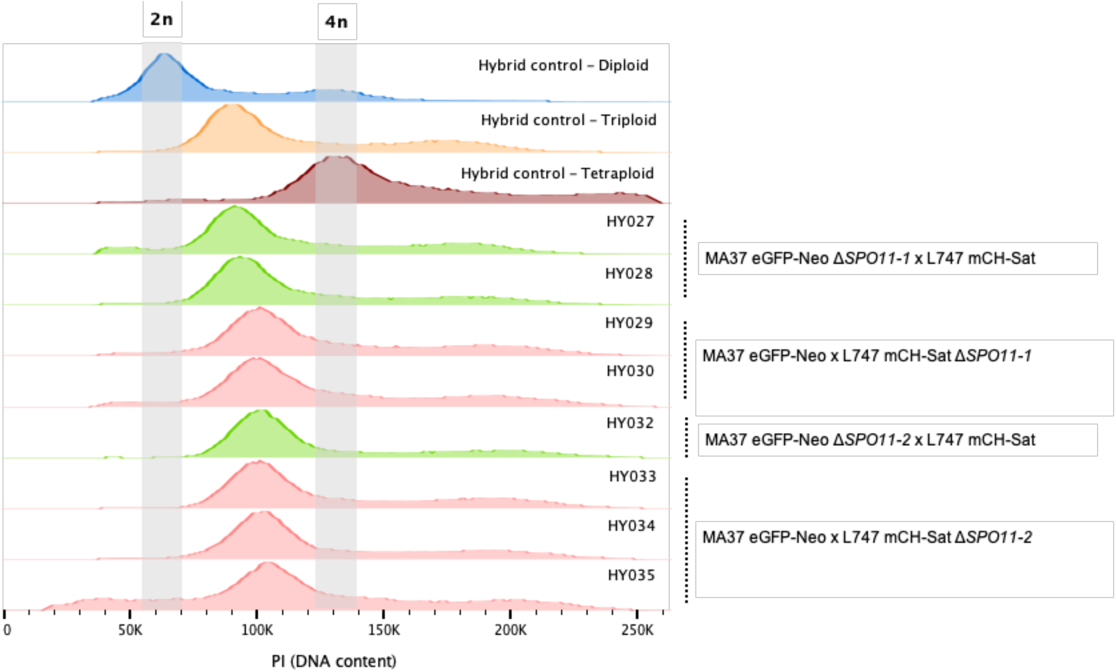

### Figure S4

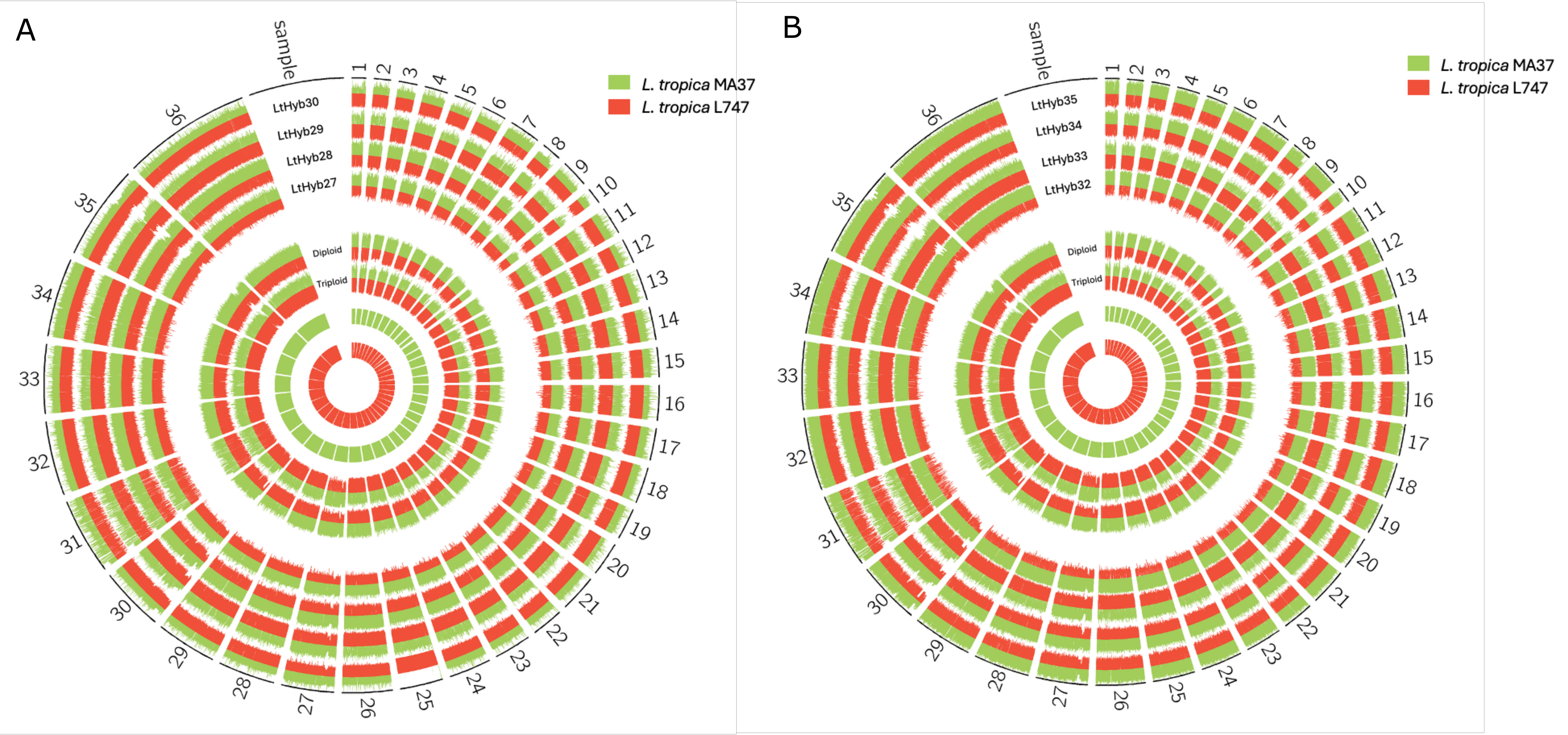
